# Spontaneous Fluctuations in Pupil Size Shape Retinal Responses to Visual Stimuli

**DOI:** 10.1101/2024.05.14.593932

**Authors:** Sebastiaan Mathôt, Daria Weiden, Olaf Dimigen

## Abstract

Visual perception is shaped at the earliest stage by the size of the eye’s pupil, which determines how much light enters the eye and how well this light is focused. But the exact role of pupil size in visual perception is still poorly understood. We recorded pupil size and electrical activity from the retina and brain while healthy human participants viewed full-screen flashes. We found that early retinal responses, which peaked ±25 ms after stimulus onset and predicted subsequent activity over visual cortex, were strongly affected by stimulus intensity. Importantly, pupil size, at least within the range of naturally occurring fluctuations, did not affect the amplitude of these early retinal responses, despite resulting in substantial changes in retinal light exposure. However, the direction of pupil-size change at the moment of stimulus presentation did modulate the amplitude of early retinal responses, which were enhanced during phases of dilation as compared to constriction. Based on these results, we suggest that fast-acting adaptation processes may normalize early retinal responses with respect to changes in retinal light exposure that result from spontaneous changes in pupil size: an initial form of brightness constancy. These results shed new light on, and raise important and previously unasked questions about, the role of pupil size in visual perception.

## Spontaneous Fluctuations in Pupil Size Shape Retinal Responses to Visual Stimuli

The size of the eyes’ pupils determines how much light enters the eye (visual sensitivity is highest for large pupils) as well as how sharply this light is focused (visual acuity is highest for small pupils; Douglas, 2018; Loewenfeld, 1958; Mathôt, 2018). However, little is currently known about the exact role that pupil size plays in visual perception, and crucial questions remain unresolved: How do changes in pupil size affect responses at different stages of the visual pathway, including the retina? How does this in turn affect behavioral performance on different tasks? And what are the functional benefits—if any—of changes in pupil size, including the small changes induced by arousal and other cognitive factors (Vilotijević & Mathôt, 2023)? To start answering these fundamental questions, here we aim to 1) empirically investigate how changes in pupil size affect retinal responses to visual stimuli, and 2) outline a new and testable hypothesis about the role of pupil size in the earliest stage of visual perception.

To what extent do changes in pupil size and changes in environmental luminance, which both affect retinal light exposure, have similar effects on visual processing? We recently used a red-blue induction technique (see also Vilotijević & Mathôt, 2024; Wardhani et al., 2022) to experimentally induce small or large pupils while recording electroencephalographic (EEG) responses to visual stimuli (Mathôt et al., 2023; for related work see Bombeke et al., 2016; Suzuki et al., 2019; Thigpen et al., 2018). We also experimentally manipulated the luminance of these stimuli. We found that both pupil size and stimulus luminance strongly affected EEG responses, mainly over occipital cortex. But crucially, the effects of pupil size and stimulus luminance, while both strong, were qualitatively distinct, such that pupil size mainly affected high-frequency (beta) activity, whereas stimulus luminance mainly affected lower frequency (alpha) activity. In other words, at least at the level of the visual cortex, the effects of changes in pupil size on visual perception are not simply equivalent to the effect of changes in retinal light exposure. However, this result does not speak to the fundamental question of whether and how pupil size already affects the *earliest* stages of visual perception—retinal responses—which cannot be picked up through EEG.

Electroretinography (ERG) is a powerful technique to study retinal responses to visual stimuli. Like EEG, ERG involves electrodes that pick up electrical activity, in this case from the retina. ERG is often used as a clinical tool to diagnose visual defects (Robson et al., 2018), in which case special electrodes are typically used that are placed directly onto the cornea; such electrodes are able to pick up retinal responses that emerge almost instantaneously and peak around 10 ms after stimulus onset; this so-called a-wave response is believed to reflect activation of rod and cone photoreceptors (Granit, 1933). Alternatively, standard skin electrodes can be placed near the eyes; such electrodes pick up responses that emerge somewhat later, but also originate from the retina (Pratt et al., 1982); this initial response as measured with standard skin electrodes likely reflects activation of various retinal neurons (including bipolar and ganglion cells), which follows photoreceptor activation. This is the response that we will focus on in the present study.

ERG studies have consistently found that ERG responses scale with stimulus intensity and pupil size, and the widespread (and intuitive) assumption seems to be that the effect of pupil size on ERG amplitude is a direct result of the fact that large pupils increase retinal light exposure (e.g. Gonzalez et al., 2004; Robson et al., 2018). However, ERG studies have generally compared very large differences in pupil size (several mm) by contrasting pharmacologically dilated pupils with either pharmacologically constricted pupils (Gonzalez et al., 2004) or natural (untreated) pupils (Mobasserian et al., 2022). Importantly, such pharmacologically induced changes in pupil size also affect subjective brightness perception (Sulutvedt et al., 2021) in a way that naturally occurring fluctuations in pupil size do not (Wardhani et al., 2022); therefore, it is unclear how relevant such large pupil-size changes are to natural visual perception. Whether smaller changes in pupil size (±1 mm), which in part reflect spontaneous fluctuations in arousal (Lowenstein et al., 1963; Mathôt et al., 2015; Reimer et al., 2014), similarly affect ERG amplitude is currently unclear. One study found that individual differences in pupil size do not correlate with individual differences in ERG amplitude (Mobasserian et al., 2022). However, to our knowledge, no study has looked at whether ERG amplitude is affected by spontaneous fluctuations in pupil size. However, from a perceptual point of view, this is a crucial question, because it relates to whether the visual system is, already at the level of the retina, able to distinguish changes in retinal light exposure due to changes in environmental brightness from self-induced changes in retinal light exposure due to differences in pupil size. In other words, it relates to whether a rudimentary form of brightness constancy with respect to pupil size is already implemented at the level of the retina.

In the present study, we simultaneously recorded pupil size and electrical activity from the retina and brain while participants viewed full-screen flashes of different intensities. Based on the literature reviewed above, we initially (and incorrectly!) hypothesized that larger pupils would lead to increased ERG amplitudes but not, or much less so, to increased ERP amplitudes over occipital cortex; in other words, we hypothesized that retinal responses would not show any brightness constancy with respect to pupil size, but reflect purely the amount of light entering the eye as a function of both stimulus intensity and pupil size, and that only responses from occipital cortex would show a degree of brightness constancy, such that they reflect mainly stimulus intensity (or contrast with the surroundings) and no longer pupil size (cf. Mathôt et al., 2023).

However, we instead found a pattern that is much more intricate and functionally adaptive than we had hypothesized. To preview the results, we found that pupil size, at least within the range of naturally occurring fluctuations, does not affect the amplitude of the earliest ERG component, despite resulting in substantial changes in retinal light exposure. However, the direction of spontaneous pupil-size change at the moment of stimulus presentation did affect the amplitude of the earliest ERG component, which was enhanced during dilation as compared to constriction.

Based on these results (see Discussion for details), we propose that fast-acting adaptation processes (Baccus & Meister, 2002; Poot et al., 1997) may normalize retinal responses, such that their amplitude is not, or hardly, affected by changes in light influx that result from spontaneous changes in pupil size: an initial form of brightness constancy with respect to pupil size that is implemented at the level of the retina. The finding that larger pupils do not lead to stronger early ERG responses seems at odds with the finding that larger pupils are associated with enhanced visual sensitivity (Ruuskanen et al., 2024). To reconcile these findings, we put forward as a testable hypothesis that increased pupil size may enhance visual sensitivity by reducing the variability, rather than modulating the strength, of early retinal responses; this would explain why, within the range of naturally occurring pupil-size fluctuations, larger pupils allow us to more easily detect faint stimuli (Ruuskanen et al., 2024) without causing us to perceive stimuli as being brighter (Wardhani et al., 2022).

## Results

Healthy human observers viewed 100 ms full-screen flashes of varying intensity (Fig. 1d). The observer’s task was to report the presence of a rare (10% of trials) target stimulus embedded in the flash stimulus. Pupil size, electrical retinal activity (henceforth electroretinogram, or *ERG*), and electrical brain activity from three occipital electrodes (O1, Oz, O2; henceforth event-related potentials, or *ERP*) were simultaneously recorded and time-locked to the onset of the flash. Unless otherwise indicated, statistical outcomes are based on linear-mixed effect, cluster-based permutation tests with stimulus intensity, pupil size, and pupil-size change as fixed effects (to limit model complexity, interaction terms were not included; adding interaction terms did not change any of the statistical outcomes, see Supplementary Results) and voltage (ERG or ERP) as dependent measure. In addition, a single linear mixed effects analysis with the same structure was performed using the peak amplitude of the first ERG component (see Methods for details).

**Figure 1.**
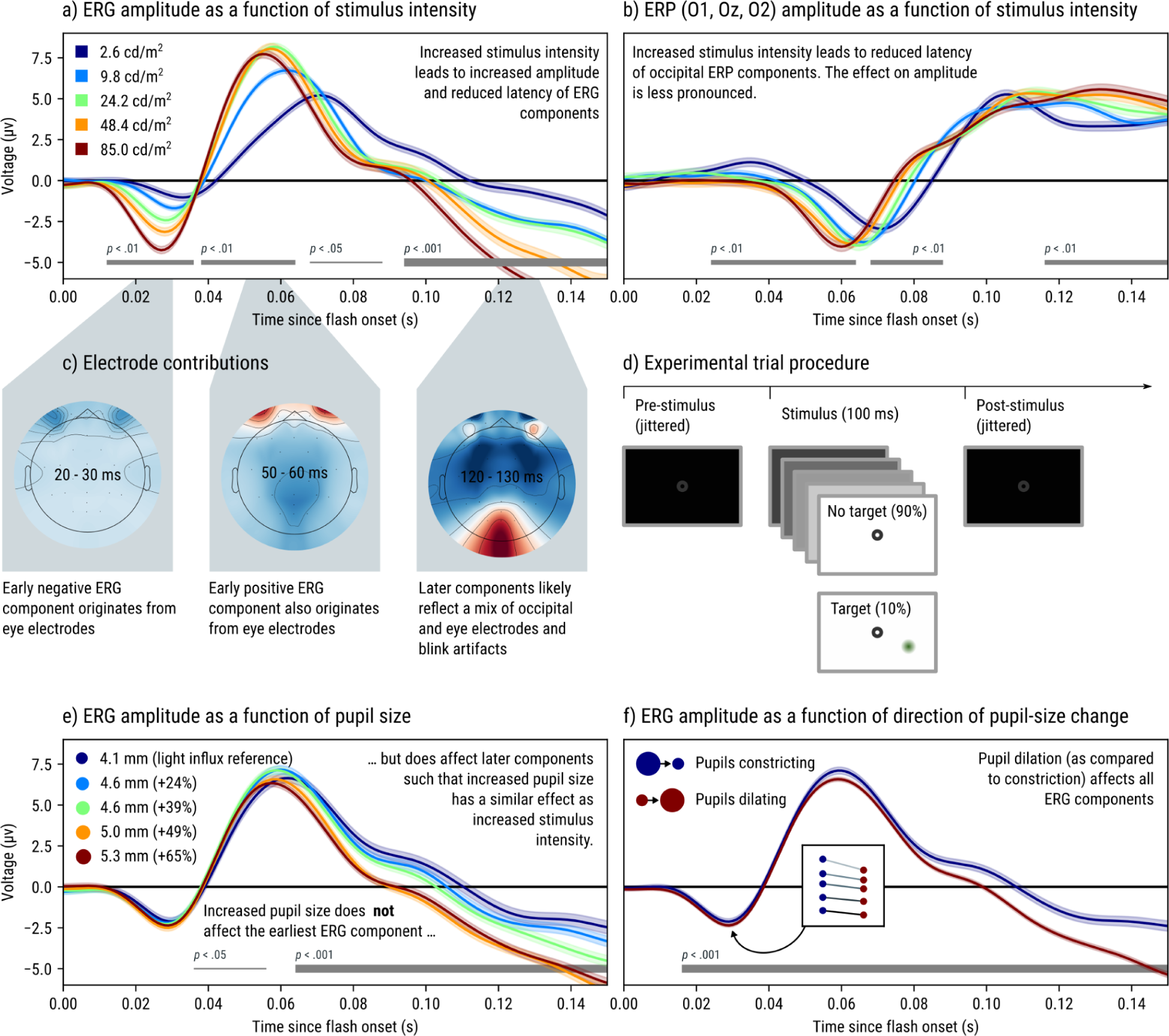
a) Stimulus intensity strongly affects ERG amplitude and latency, such that components are both more pronounced and peak faster for high-intensity as compared to low-intensity stimuli. b) Stimulus intensity mainly affects occipital (mean across O1, Oz, O2) ERP latency, such that the component peaks faster for high-as compared to low-intensity stimuli; differences in ERP amplitude are less pronounced. c) Scalp topographies including all electrodes (EEG and ERG) show that early activity originates purely from the eye electrodes, whereas later activity is more distributed, originates also from occipital cortex, and likely also contains artifacts related to (partial) eye blinks (see Supplementary Results). d) Each trial started with a dark display with a fixation dot. This was followed by a 100 ms flash of varying levels of intensity. On 10% of trials, a target stimulus (a green patch with a gaussian envelope) was embedded in the flash stimulus. Participants were instructed to press the spacebar whenever they saw a target. e) Spontaneous fluctuations in pupil size did not affect the earliest ERG component, despite changing the amount of light influx considerably. Later ERG components were affected by pupil size, such that the effect of larger pupils at later time points (> 60 ms) was qualitatively similar to the effect of increased stimulus intensity. Percentages in the legend indicate the increase in light influx relative to the bin of smallest pupil sizes. f) Pupil-size change affected ERG amplitude from the very first response (< 20 ms after flash onset). The inset panel indicates peak amplitude of the first component as a function of intensity (separate lines) and pupil-size change. a, b, e, f) Gray horizontal lines indicate clusters in which there was a significant effect of stimulus intensity on voltage (thin: p < .05, medium: p < .01, thick: p < .001). Error bands indicate standard error of the grand mean.

### General ERG, ERP, and eye-blink characteristics

The first ERG component emerged as a negative deflection around 10 ms after stimulus onset and peaked around 25 ms for the highest intensity level (Fig. 1a). This was followed by a second, positive component that peaked around 50-70 ms (depending on stimulus intensity, see below). These first two ERG components originated entirely from the eye electrodes (Fig. 1c), with the same polarity above and below the eye (relative to the reference electrodes located over the left and right mastoid process), suggesting that they reflect retinal activity (Pratt et al., 1982) rather than eye movements or blinks. We do not attempt to interpret these components in terms of a-wave and b-wave ERG components, which are generally obtained using Ganzfeld stimulation and corneal electrodes (Robson et al., 2018).

The first ERP component, likely reflecting the C1 component of visually evoked potentials (e.g. Clark et al., 1994), emerged as a negative deflection around 40 ms and peaked at around 50-70 ms over midline occipital electrodes. Granger causality analysis suggested that ERG activity preceded and predicted occipital ERP activity (see Supplementary Results for details), consistent with a flow of information from the retina to the brain.

Since we presented stimuli on an LCD monitor, response latencies in part reflect the properties of the monitor (e.g. Kaltwasser et al., 2009; Keating et al., 2001). Specifically, it takes some time for the entire LCD monitor to refresh and for individual pixels to reach maximum brightness, both of which result in a degree of temporal smearing. Here, response latencies are measured from the moment that pixels on the top-left corner of the display started to increase in brightness (see Supplementary Results).

Participants frequently blinked in response to the flash stimulus. Although blinks typically only occurred about 200 ms after flash onset, the ERG signal was modulated by the presence of blinks already from around 60 ms (see Supplementary Results for details). Although we excluded trials in which a blink occurred within 500 ms after stimulus onset (see Methods), the signal appeared to still be contaminated by blink-like artefacts, likely caused by partial closures of the eyelid. For this reason, our analysis and conclusions are focused on the earliest ERG component, which remained entirely uncontaminated by blinks.

### Effects of stimulus intensity on ERG and ERP

#### ERG: stimulus intensity affects amplitude and latency (Fig. 1a)

Increased stimulus intensity strongly affected the ERG from 12 ms onwards (onset of the first component) in a more-or-less contiguous sequence of statistical clusters until the end of the analysis period (12 - 36 ms: *p* = .003; 38 - 64 ms: *p* = .002; 86 - 88 ms: *p* = .025, 94 - 150 ms; *p* < .001). The amplitude of components increased with increased stimulus intensity, whereas latency decreased. Qualitatively, the ERG response to a weak stimulus was reduced, temporally delayed, and temporally smeared as compared to the response to a strong stimulus.

#### ERP: stimulus intensity mainly affects latency (Fig. 1b)

Increased stimulus intensity affected the occipital ERP from 24 ms onwards in a series of clusters (24 - 64 ms: *p* = .001; 68 - 88 ms: *p* = .005; 116 - 150 ms: *p* = .001), a period roughly coinciding with the C1 component (note that the extremely early start of the first C1 cluster [24 ms] likely reflects the tendency of cluster-based permutation tests to underestimate onset latencies; Sassenhagen & Draschkow, 2019). Qualitatively, stimulus intensity mainly affected the latency of the C1, whereas stimulus intensity had only a minor influence on C1 amplitude.

### Effects of pupil size and pupil-size change (dilation vs constriction) on ERG

For each trial, we took the mean pupil size during the first 150 ms after stimulus onset as our measure of pupil size. This period was chosen to 1) fully cover the flash duration, 2) be long enough to estimate pupil size as well as the direction of pupil-size change, and 3) short enough to avoid the pupil-constriction response to the flash stimulus, which has a latency of about 200 ms (see Supplementary Results), from affecting our pupil measures. In addition, we fitted a linear regression through the pupil-size time series during this 150 ms window, and took the regression slope as our measure of pupil-size change. Trials with a positive pupil-size change were considered Dilating trials, whereas trials with a negative or zero slope were considered Constricting trials. For visualization we divided trials into five pupil-size bins with an equal number of trials (for pupil-size analyses) or Dilating and Constricting trials (for pupil-size-change analyses); however, statistical analyses were performed on pupil size or pupil-size change as continuous measures.

Importantly, our pupil measures are distinct from the pupil-constriction response to the stimulus, which occurs later. Rather, our measures reflect the state of the pupil during the moment that the stimulus was presented.

In our study, changes in pupil size were the result of naturally occurring fluctuations, rather than an experimental manipulation. Because we assume that effects of pupil size on ERG responses are primarily related to processes that happen locally in the retina, we phrase correlations between pupil size and ERG activity as effects of pupil size on ERG activity. However, in the discussion we will return to possible contributions of (retinopetal) feedback connections from the brain to the retina (Gastinger et al., 2006).

#### ERG: pupil size does not affect the earliest component, but does affect later components (Fig. 2e)

Strikingly, pupil size did not affect the earliest ERG component that emerged around 10 ms after flash onset. This is remarkable because this component is highly sensitive to changes in stimulus intensity, and naturally occurring pupil-size changes are large enough to result in substantial changes in light influx: the 20% largest pupils resulted in a 65% increase in light influx relative to the 20% smallest pupils.

This null result was reflected in the lack of a statistical cluster that spanned the earliest ERG component in the cluster-based permutation test, as well as the lack of an effect of pupil size on the amplitude of the first ERG component in the single linear mixed-effects model (*z* = -0.93, *p* = .352). To substantiate this null result further, we compared two linear mixed-effects models: an “intensity model” in which the amplitude of the first ERG component was predicted only by stimulus intensity, and a “light-influx model” in which the amplitude was predicted by stimulus intensity as well as the interaction between stimulus intensity and an adjustment factor that, for each trial, up- or down-regulated the effect of stimulus intensity based on pupil size. (For example, for a trial on which the pupil let in 10% more light than the average trial for a given participant, the adjustment factor would be 1.1.) Model comparison revealed that the intensity model (AIC = -98914) was a slightly better fit to the data than the light-influx model (AIC = -98912), confirming that within the range of pupil sizes that we observed here, pupil size did not notably affect the earliest component.

However, pupil size did affect later ERG components, starting around 36 ms after stimulus flash onset and, in two clusters, more or less continuously until the end of the analysis period (36 - 56 ms: *p* = .033; 64 - 150 ms: *p* < .001). Qualitatively, the effect was such that, at these later time points, an increase in pupil size had a similar effect on ERG activity as an increase in stimulus intensity.

#### ERG: pupil-size change (dilation vs constriction) affects all ERG components (**Fig. 2f**)

Pupil-size change affected ERG amplitude from 16 ms after stimulus onset (roughly corresponding to the onset of the first component) until the end of the analysis period (16 - 150 ms; *p* < .001) in the cluster-based permutation test. Pupil-size change also affected the amplitude of the first ERG component in the single linear-effects model (*z* = -3.01, *p* = .003). This was a unique effect of pupil-size change, and not driven by correlations with absolute pupil size (because both were included as predictors in the same model) nor by carry-over effects from the previous trial (see Supplementary Results). Qualitatively, the effect was such that an increase in pupil dilation had a similar effect on ERG activity as an increase in stimulus intensity.

## Discussion

Here we report two key findings related to the effect of pupil size on retinal responses to visual stimuli. 1) Natural fluctuations in pupil size do not notably affect the amplitude of the first electroretinographic (ERG) component that we were able to pick up with our methodology, which emerges around 10 ms after stimulus onset, peaks around 25 ms, and originates exclusively from eye electrodes. This is a crucial null-result, because such pupil-size fluctuations result in substantial differences in retinal light exposure (>60%), which when manipulated by varying stimulus intensity, strongly affect this early ERG component. 2) A reliable predictor of the amplitude of the first ERG component is whether, at the moment that the stimulus appears, the pupil is dilating (becoming larger) or constricting (becoming smaller), regardless of the absolute size of the pupil.

One interpretation of our results, which at present we favor because of its simplicity, requires only retinal adaptation; more specifically, it requires a form of fast-acting retinal adaptation that takes place on a time scale of hundreds of milliseconds (Baccus & Meister, 2002; Poot et al., 1997). (Such fast-acting adaptation processes occur in addition to much slower adaptation processes that occur on a time scale of seconds or minutes [Meister & Berry, 1999] and that likely do not play a role in our results.)

To understand how this could explain our observations, consider a dimly lit (but not fully dark) lab environment in which participants are observing a static (non-flashing) display. When pupils are large, and thus allow more light to reach the retina, retinal adaptation is slightly stronger than when pupils are small, and thus allow less light to reach the retina. Now consider a flash that is briefly presented on this display. When pupils are large as compared to small, the retina is exposed to more light from this flash, which increases the strength of the retinal response; however, the retina is also in a slightly stronger state of adaptation, which decreases the strength of the response; these two counteracting forces cancel each other out, in our case even almost perfectly. As a result, the amplitude of the first ERG component becomes invariant to changes in pupil size, at least within the range of naturally occurring fluctuations and within the range of stimulus intensities tested here (see Gonzalez et al., 2004; Mobasserian et al., 2022 for studies using pharmacological manipulations of pupil size). Functionally, this may be one of many processes that contribute to brightness constancy, and more specifically brightness constancy with respect to natural fluctuations in pupil size.

Now consider a flash that is presented while the pupil is dilating. In this case, the retina is in a slightly reduced state of adaptation (because a dilating pupil was by definition smaller in the recent past), yet receives increased light influx (because the pupil is now larger than it was moments before). As a result, the amplitude of the retinal response increases; more generally, the amplitude of the retinal response becomes modulated by whether the pupil is dilating or constricting at the moment that a stimulus is presented, which is precisely what we observed. There is no clear functional interpretation of the finding that early ERG amplitude is modulated by the direction of pupil-size change. Tentatively, however, it may be related to studies showing that pupil dilation (more so than absolute pupil size) is in some cases associated with increased behavioral performance (Brink et al., 2016) and bursts of neural activity (Reimer et al., 2014). In addition, a prediction that follows from this finding is that the perceived brightness of a stimulus should be affected by the direction of pupil-size change at the moment of stimulus presentation, such that pupil dilation is associated with increased perceived brightness.

An alternative interpretation of our results involves feedback from the brain through so-called retinopetal axons that project from the hypothalamus to the retina (Gastinger et al., 2006). Although such axons are few in number and their function is poorly understood, there is preliminary evidence that direct brain-to-retina feedback may allow higher-level cognitive processes to directly modulate visual processing at the level of the retina (Schröder et al., 2020; Warwick et al., 2022). This is clearly an exciting possibility, but, as discussed above, the present results can likely be explained through known mechanisms and without reference to (speculative) direct brain-to-retina feedback.

The absence of an effect of pupil size on the strength of early retinal responses seems at odds with the well-established finding that larger pupils are associated with increased visual sensitivity (Eberhardt et al., 2022; Mathôt & Ivanov, 2019). For example, Ruuskanen et al. (2024) recently found that larger pupils, as in the current study resulting from natural fluctuations, are associated with enhanced visual sensitivity in a near-threshold visual detection task; importantly, a mediation analysis indicated that this association reflects, at least in part, a direct effect of pupil size on sensitivity (as opposed to an indirect effect mediated by various EEG markers of arousal and cortical excitability that were considered in this study). But if pupil size does not affect early retinal responses, then how can pupil size affect visual sensitivity at a behavioral level?

We conjecture that larger pupils may be associated with a reduced variability of retinal activity; phrased differently, larger pupils may make the response to a stimulus more consistent, rather than making the response stronger. Most models of neural coding, including of the retina (Cui et al., 2016; Meister & Berry, 1999), implement some form of divisive normalization (Carandini & Heeger, 2012). If retinal adaptation also approximates a divisive operation, this would mean that the strength of the retinal response is proportional to retinal light exposure divided by adaptation state. As a result, increased adaptation would reduce both the strength and variability of the response, which in turn may lead to improved visual sensitivity. The conjecture that larger pupils lead to reduced variability in the earliest ERG response is testable in future studies, provided that there is a clear definition of the specific measure of variability (or information or decoding) that is relevant in this context (Quian Quiroga & Panzeri, 2009), as well as a large dataset that allows for precise estimates of variability.

Alternatively, the fact that larger pupils are associated with improved visual sensitivity at a behavioral level may be related to later stages of retinal activity; specifically, even though the amplitude of the first ERG component was not affected by pupil size, ERG amplitude at later points in time was associated with pupil size (especially >60 ms after stimulus onset). However, at these later points in time, the ERG signal is contaminated with artifacts due to eyelid movements and potentially also by cortical potentials that reach the eye electrodes via volume conduction. Because of this, it is difficult to interpret the association between pupil size and later stages of the ERG signal; for this reason, we have largely limited our discussion to the first ERG component, which purely reflects retinal activity.

In summary, we found that natural fluctuations in pupil size do not affect the strength of early retinal responses, despite resulting in substantial changes in retinal light exposure; however, the direction of pupil-size change at the moment of stimulus presentation does modulate the strength of early retinal responses, which are enhanced during pupil dilation as compared to constriction. We propose that these results can be explained through fast-acting retinal adaptation processes. Taken together, these results suggest that an initial form of brightness constancy with respect to pupil size is already implemented at the level of the retina, and raise important and previously unasked questions about the role of pupil size in the earliest stages of visual perception.

## Methods

### Open-practices statement

All data, experimental materials, and analysis scripts are available from https://osf.io/g5utq/.

The study was not preregistered, and consists of confirmatory (corresponding to Fig. 1a-c, e) and exploratory (Fig. 1f) analyses. We will therefore emphasize general patterns over significance of individual effects. When reporting statistical analyses, we will refer to *p* < .05 as ‘significant effects’ (i.e., using an alpha level of .05). All analyses are corrected for multiple comparisons.

### Participants

10 participants (including authors SM and DW) participated in the experiment. The sample size was predetermined based on pilot studies, which showed that early ERG responses are highly replicable between participants as well as between different sessions of the same participant; based on this consideration, combined with the fact that participants may feel uncomfortable having electrodes attached so close to the eye, we decided to extensively test a small sample of participants that felt comfortable with the procedure. One additional participant was tested but excluded from analysis and replaced because of ERG responses that, based on visual inspection, strongly deviated from those of the other participants due to frequent blinking.

### Materials and software

The experimental script was implemented using OpenSesame 4.0 (Mathôt et al., 2012) using the PsychoPy (Peirce, 2007) backend for display presentation and PyGaze (Dalmaijer et al., 2014) for eye tracking.

The experiment was conducted on a desktop computer with a 27” LCD monitor with a refresh rate of 120 Hz and a resolution of 1920 × 1080 pixels. At the viewing distance of 76 cm, the full-screen stimuli subtended 44.8° x 25.2° degrees of visual angle. Pupil diameter and gaze position were recorded from the right eye at 1000 Hz using an EyeLink 1000 (SR Research) eye tracker. EEG data was recorded at 1000 Hz from 26 electrodes placed according to the standard 10-20 system using a TMSI REFA 32 amplifier controlled by OpenViBE data acquisition software (Renard et al., 2010). Two additional electrodes were placed on the left and right mastoids for offline re-referencing. Two additional channels were placed immediately above the eyelids and on the infraorbital margin of each eye (four channels in total; Fig. 3). During recording all electrodes (EEG and ERG) were referenced against the average of all electrodes, and then re-referenced offline against the average of the left and right mastoids.

**Figure 3.**
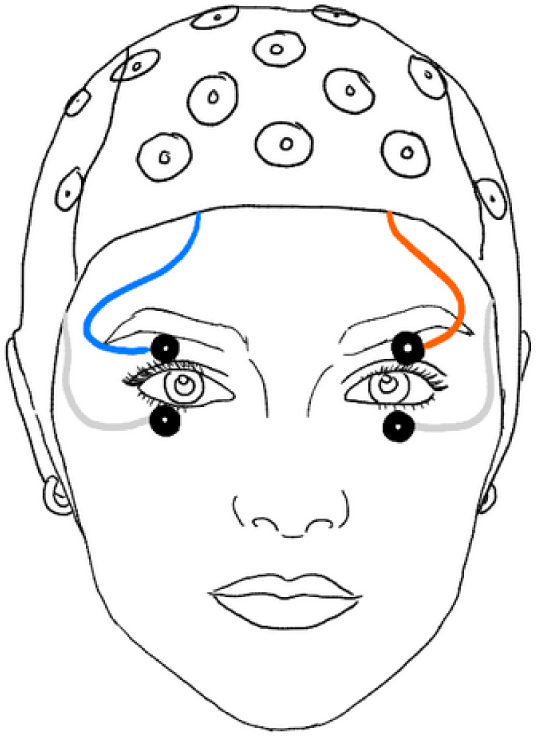
EOG electrodes were placed above and below each eye. The ERG signal as shown in Fig. 1 and 2 correspond to the average of those four channels.

Data analysis was performed using the Python `eeg_eyetracking_parser` (Mathôt et al., 2023) toolbox, which is a high-level toolbox that uses `MNE` (Gramfort et al., 2013) for general EEG processing, `eyelinkparser` (Mathôt & Vilotijević, 2022) for eye-movement and pupil-size processing, and `autoreject` (Jas et al., 2017) for rejecting and repairing bad EEG channels and epochs. Cluster-based permutation tests were performed using `time_series_test` (Mathôt & Vilotijević, 2022).

### Procedure

The experiment took place in a dimly lit room (<1 lux). This corresponds to mesopic vision: lighting conditions in which vision is mediated by both rod and cone photoreceptors, and in which dark adaptation has not taken place (Stockman & Sharpe, 2006).

Each participant completed two sessions of 500 trials each, which were in turn divided into blocks of 100 trials. Each session started with a five-point eye-tracking calibration procedure, followed by two instruction screens. Each session was a separate recording, but all participants opted to do both sessions on the same day.

Each trial started with a small, dim fixation point centered on a dark (<0.01 cd/m²) background. Next, a full-screen flash was presented for 100 ms. The intensity of this flash varied in steps that were perceptually linear in CIELAB color space (2.69 cd/m², 9.84 cd/m², 24.24 cd/m², 48.45 cd/m², 85 cd/m²). On 10% of trials, a target (a green patch with a gaussian envelope [SD = 0.77°] and a contrast proportional to the intensity of the flash) was presented at a random location anywhere on the display, followed by an interval jittered between 1500 and 2000 ms during which participants could press a key to report the presence of the target; this was an easy task that resulted in few misses or false alarms, and simply served to ensure that participant kept attending to the flashes. Response accuracy was therefore not further analyzed. Unobtrusive feedback was provided after the response interval by briefly presenting a green fixation dot after a hit, and a red fixation dot after a false alarm or a miss; no feedback was provided after a correct rejection. The median interval between flashes was 3317 ms.

### Data preprocessing

#### Pupil size

For the pupil-size analyses, we followed the recommendations from Mathôt and Vilotijević (2022). 1) Missing or invalid data was interpolated using cubic-spline interpolation if possible, using linear interpolation if cubic-spline interpolation was not possible (when the segment of missing data was too close to the start or end of a trial), and removed if interpolation was impossible altogether (when data was missing from the start and/ or until the end of a trial) or if the period of missing data was longer than 500 ms and thus unlikely to reflect a blink (see `datamatrix.series.blinkreconstruct`). 2) Pupil size was converted from arbitrary units as recorded by the EyeLink to millimeters of diameter. No baseline correction was applied to pupil size.

#### Electroencephalography (EEG) and electroretinography (ERG)

EEG and ERG data was preprocessed fully automatically. Unless otherwise specified, we used the default parameters as described on the documentation of the referenced functions. 1) Data was re-referenced to the average of the mastoid channels. 2) Muscle artifacts, which are characterized by bursts of high-frequency activity, were marked as bad using the MNE function for annotating muscle activity (see `mne.preprocessing.annotate_muscle_zscore`) with a z-threshold of 5. 3) Data was filtered using a 0.1 - 40 Hz bandpass. 4) The RANSAC algorithm (see `autoreject.Ransac`) identified bad channels based only on data segments corresponding to trials; in brief, this algorithm assumes that a channel is bad if its data is poorly predicted by interpolation from neighboring channels (Bigdely-Shamlo et al., 2015). Baseline correction was applied to the ERG or EEG signal (but not to pupil size) using the 100 ms preceding stimulus onset as the baseline period.

For analyses where ERG amplitude was the dependent variable (Fig. 1a. and Fig 2), the signal was based on the average of all four EOG channels, referenced against the mean of the left and right mastoid. For analyses where EEG amplitude was the dependent variable (Fig. 1b), the signal was defined as the average of the three occipital electrodes (O1, Oz, and O2), also referenced against the mean of the left and right mastoid.

#### Trial exclusion

Trials were excluded from analysis if 1) pupil-size data was missing for the full 150 ms after stimulus onset, 2) mean pupil size (averaged across the 150 ms after stimulus onset), mean pupil-size change (slope), or the the average amplitude between 15 and 40 ms post-stimulus deviated more than 3 SD from the participant’s mean value for those respective measures, 3) a blink as identified by the EyeLink’s built-in algorithm occurred within 500 ms after stimulus onset, or 4) a target was presented on that trial. After exclusion, 7749 trials (77.5%) remained for further analysis.

### Statistical analysis

Our analyses are based on single-trial, linear-mixed effect, cluster-based permutation tests with stimulus intensity, pupil size, and pupil-size change as fixed effects and voltage as dependent measure (see `time_series_test.lmer_permutation_test`). Fixed effects (but not random effects or dependent measures) were z-scored to improve convergence. This analysis was conducted separately for ERG and EEG voltage. Models included by-participant random intercepts, but no by-participant random slopes; this choice was motivated by our pilot data, which revealed limited individual differences in the ERG responses. Interaction terms were not included to limit model complexity. Clusters were identified using a *p* < .05 criterion, and the absolute sum of z-scores for individual clusters was used as test statistic. We used windows of 2 ms and ran 1,000 iterations. Significant clusters provide only a rough indicator of when effects arise (Sassenhagen & Draschkow, 2019). Nevertheless, we also report the cluster extents here, since they provide a useful indication of the temporal extent of the effects and also coincide with periods in which effects are clearly visible in the waveforms.

In addition, we conducted a single linear-mixed effects model that used the peak of the first ERG component as dependent measure, and that otherwise used the same approach as the cluster-based permutation test. To estimate the (negative) peak of the first ERG component, we took the sample with the lowest voltage in the 15 - 40 ms window in the mean signal per participant and intensity level; splitting by participant and intensity level was done to account for the fact that there are likely differences in the latency of the component between participants and intensity levels. For this analysis, trials with a peak amplitude that deviated more than 2 SD from the mean peak amplitude were excluded (4.1%). This decision was based on a visual inspection of the distribution of peak amplitudes; including all trials did not change any of the statistical outcomes.

## Supporting information

Supplementary Results

## References

Baccus, S. A., & Meister, M. (2002). Fast and slow contrast adaptation in retinal circuitry. Neuron, 36(5), 909–919. 10.1016/S0896-6273(02)01050-4

Bigdely-Shamlo, N., Mullen, T., Kothe, C., Su, K.-M., & Robbins, K. A. (2015). The PREP pipeline: Standardized preprocessing for large-scale EEG analysis. Frontiers in Neuroinformatics, 9. 10.3389/fninf.2015.00016

Bombeke, K., Duthoo, W., Mueller, S. C., Hopf, J.-M., & Boehler, N. C. (2016). Pupil size directly modulates the feedforward response in human primary visual cortex independently of attention. NeuroImage, 127, 67–73. 10.1016/j.neuroimage.2015.11.072

Brink, R. L. van den, Murphy, P. R., & Nieuwenhuis, S. (2016). Pupil diameter tracks lapses of attention. PLOS ONE, 11(10), e0165274. 10.1371/journal.pone.0165274

Carandini, M., & Heeger, D. J. (2012). Normalization as a canonical neural computation. Nature Reviews Neuroscience, 13(1), 51–62. 10.1038/nrn3136

Clark, V. P., Fan, S., & Hillyard, S. A. (1994). Identification of early visual evoked potential generators by retinotopic and topographic analyses. Human Brain Mapping, 2(3), 170–187. 10.1002/hbm.460020306

Cui, Y., Wang, Y. V., Park, S. J. H., Demb, J. B., & Butts, D. A. (2016). Divisive suppression explains high-precision firing and contrast adaptation in retinal ganglion cells. eLife, 5, e19460. 10.7554/eLife.19460

Dalmaijer, E., Mathôt, S., & Van der Stigchel, S. (2014). PyGaze: An open-source, cross-platform toolbox for minimal-effort programming of eyetracking experiments. Behavior Research Methods, 46(4), 913–921. 10.3758/s13428-013-0422-2

Douglas, R. H. (2018). The pupillary light responses of animals; a review of their distribution, dynamics, mechanisms and functions. Progress in Retinal and Eye Research, 66, 17–48. 10.1016/j.preteyeres.2018.04.005

Eberhardt, L. V., Strauch, C., Hartmann, T. S., & Huckauf, A. (2022). Increasing pupil size is associated with improved detection performance in the periphery. *Attention, Perception*, & Psychophysics, 84(1), 138–149. 10.3758/s13414-021-02388-w

Gastinger, M. J., Tian, N., Horvath, T., & Marshak, D. W. (2006). Retinopetal axons in mammals: Emphasis on histamine and serotonin. Current Eye Research, 31(7–8), 655–667. 10.1080/02713680600776119

Gonzalez, P., Parks, S., Dolan, F., & Keating, D. (2004). The effects of pupil size on the multifocal electroretinogram. Documenta Ophthalmologica, 109(1), 67–72. 10.1007/s10633-004-1545-7

Gramfort, A., Luessi, M., Larson, E., Engemann, D., Strohmeier, D., Brodbeck, C., Goj, R., Jas, M., Brooks, T., Parkkonen, L., & Hämäläinen, M. (2013). MEG and EEG data analysis with MNE-Python. Frontiers in Neuroscience, 7. 10.3389/fnins.2013.00267

Granit, R. (1933). The components of the retinal action potential in mammals and their relation to the discharge in the optic nerve. The Journal of Physiology, 77(3), 207–239. 10.1113/jphysiol.1933.sp002964

Jas, M., Engemann, D. A., Bekhti, Y., Raimondo, F., & Gramfort, A. (2017). Autoreject: Automated artifact rejection for MEG and EEG data. NeuroImage, 159, 417–429. 10.1016/j.neuroimage.2017.06.030

Kaltwasser, C., Horn, F. K., Kremers, J., & Juenemann, A. (2009). A comparison of the suitability of cathode ray tube (CRT) and liquid crystal display (LCD) monitors as visual stimulators in mfERG diagnostics. Documenta Ophthalmologica, 118, 179–189. 10.1007/s10633-008-9152-7

Keating, D., Parks, S., Malloch, C., & Evans, A. (2001). A comparison of CRT and digital stimulus delivery methods in the multifocal ERG. Documenta Ophthalmologica, 102, 95–114. 10.1023/A:1017527006572

Loewenfeld, I. E. (1958). Mechanisms of reflex dilatation of the pupil. Documenta Ophthalmologica, 12(1), 185–448. 10.1007/BF00913471

Lowenstein, O., Feinberg, R., & Loewenfeld, I. E. (1963). Pupillary movements during acute and chronic fatigue: A new test for the objective evaluation of tiredness. Investigative Ophthalmology & Visual Science, 2(2), 138–157.

Mathôt, S. (2018). Pupillometry: Psychology, physiology, and function. Journal of Cognition, 1(1), 1–16. 10.5334/joc.18

Mathôt, S., Berberyan, H., Büchel, P., Ruuskanen, V., Vilotijević, A., & Kruijne, W. (2023). Effects of pupil size as manipulated through ipRGC activation on visual processing. NeuroImage, 283, 120420. 10.1016/j.neuroimage.2023.120420

Mathôt, S., & Ivanov, Y. (2019). The effect of pupil size and peripheral brightness on detection and discrimination performance. PeerJ, 7, e8220. 10.7717/peerj.8220

Mathôt, S., Schreij, D., & Theeuwes, J. (2012). OpenSesame: An open-source, graphical experiment builder for the social sciences. Behavior Research Methods, 44(2), 314–324. 10.3758/s13428-011-0168-7

Mathôt, S., Siebold, A., Donk, M., & Vitu, F. (2015). Large pupils predict goal-driven eye movements. Journal of Experimental Psychology: General, 144(3), 513–521. 10.1037/a0039168

Mathôt, S., & Vilotijević, A. (2022). Methods in cognitive pupillometry: Design, preprocessing, and statistical analysis. Behavior Research Methods, 55, 3055–3077. 10.3758/s13428-022-01957-7

Meister, M., & Berry, M. J. (1999). The neural code of the retina. Neuron, 22(3), 435–450. 10.1016/S0896-6273(00)80700-X

Mobasserian, A., Zaidi, M., Halim, S., Hwang, J. J., Regenold, J., Akhavanrezayat, A., Karaca, I., Khojasteh Jafari, H., Yavari, N., Matsumiya, W., Yasar, C., Than, N. T. T., Uludag, G., Do, D., Ghoraba, H., & Nguyen, Q. D. (2022). Effect of pupil size on fixed-luminance flicker full-field electroretinogram magnitude. Clinical Ophthalmology, 16, 3733–3740. 10.2147/OPTH.S382207

Peirce, J. W. (2007). PsychoPy: Psychophysics software in Python. Journal of Neuroscience Methods, 162(1–2), 8–13. 10.1016/j.jneumeth.2006.11.017

Poot, L., Snippe, H. P., & van Hateren, J. H. (1997). Dynamics of adaptation at high luminances: Adaptation is faster after luminance decrements than after luminance increments. Journal of the Optical Society of America A, 14(9), 2499–2508. 10.1364/JOSAA.14.002499

Pratt, H., Bleich, N., & Berliner, E. (1982). Short latency visual evoked potentials in man. Electroencephalography and Clinical Neurophysiology, 54(1), 55–62. 10.1016/0013-4694(82)90231-0

Quian Quiroga, R., & Panzeri, S. (2009). Extracting information from neuronal populations: Information theory and decoding approaches. Nature Reviews Neuroscience, 10(3), 173–185. 10.1038/nrn2578

Reimer, J., Froudarakis, E., Cadwell, C. R., Yatsenko, D., Denfield, G. H., & Tolias, A. S. (2014). Pupil fluctuations track fast switching of cortical states during quiet wakefulness. Neuron, 84(2), 355–362. 10.1016/j.neuron.2014.09.033

Renard, Y., Lotte, F., Gibert, G., Congedo, M., Maby, E., Delannoy, V., Bertrand, O., & Lécuyer, A. (2010). Openvibe: An open-source software platform to design, test, and use brain–computer interfaces in real and virtual environments. Presence, 19(1), 35–53. 10.1162/pres.19.1.35

Robson, A. G., Nilsson, J., Li, S., Jalali, S., Fulton, A. B., Tormene, A. P., Holder, G. E., & Brodie, S. E. (2018). ISCEV guide to visual electrodiagnostic procedures. Documenta Ophthalmologica, 136(1), 1–26. 10.1007/s10633-017-9621-y

Ruuskanen, V., Boehler, C. N., & Mathôt, S. (2024). The interplay of spontaneous pupil-size fluctuations and EEG power in near-threshold detection. bioRxiv. 10.1101/2024.05.13.593918

Sassenhagen, J., & Draschkow, D. (2019). Cluster-based permutation tests of MEG/EEG data do not establish significance of effect latency or location. Psychophysiology, 56(6), e13335. 10.1111/psyp.13335

Schröder, S., Steinmetz, N. A., Krumin, M., Pachitariu, M., Rizzi, M., Lagnado, L., Harris, K. D., & Carandini, M. (2020). Arousal modulates retinal output. Neuron, 107(3), 487–495.e9. 10.1016/j.neuron.2020.04.026

Stockman, A., & Sharpe, L. T. (2006). Into the twilight zone: The complexities of mesopic vision and luminous efficiency. Ophthalmic and Physiological Optics, 26(3), 225–239. 10.1111/j.1475-1313.2006.00325.x

Sulutvedt, U., Zavagno, D., Lubell, J., Leknes, S., Sigrid, A., & Laeng, B. (2021). Brightness perception changes related to pupil size. Vision Research, 178, 41–47. 10.1016/j.visres.2020.09.004

Suzuki, Y., Minami, T., & Nakauchi, S. (2019). Pupil Constriction in the Glare Illusion Modulates the Steady-State Visual Evoked Potentials. Neuroscience, 416, 221–228. 10.1016/j.neuroscience.2019.08.003

Thigpen, N. N., Bradley, M. M., & Keil, A. (2018). Assessing the relationship between pupil diameter and visuocortical activity. Journal of Vision, 18(6), 7–7. 10.1167/18.6.7

Vilotijević, A., & Mathôt, S. (2023). Functional benefits of cognitively driven pupil-size changes. WIREs Cognitive Science. 10.1002/wcs.1672

Vilotijević, A., & Mathôt, S. (2024). Non-image forming vision as measured through ipRGC-mediated pupil constriction is not modulated by covert visual attention. Cerebral Cortex. 10.1101/2023.06.27.546729

Wardhani, I. K., Boehler, C. N., & Mathôt, S. (2022). The influence of pupil responses on subjective brightness perception. Perception, 51(6), 370–387. 10.1177/03010066221094757

Warwick, R. A., Riccitelli, S., Heukamp, A. S., Yaakov, H., Ankri, L., Mayzel, J., Gilead, N., Parness-Yossifon, R., & Rivlin-Etzion, M. (2022). Top-down modulation of the retinal code via histaminergic neurons of the hypothalamus (p. 2022.04.26.489509). bioRxiv. 10.1101/2022.04.26.489509

