## Supplementary Results for "Spontaneous Fluctuations in Pupil Size Shape Retinal Responses to Visual Stimuli"

### Open-practices statement

All data, experimental materials, and analysis scripts are available from https://osf.io/g5utq/.

### The flash triggers a typical pupil light response

We observed a typical pupil light response to the flash stimulus (Fig. 1). The pupil constricted in response to the flash with an onset latency of around 200 ms. The strength of pupil constriction was strongly modulated by the intensity of the flash, such that the pupil constricted more strongly in response to more intense flashes.

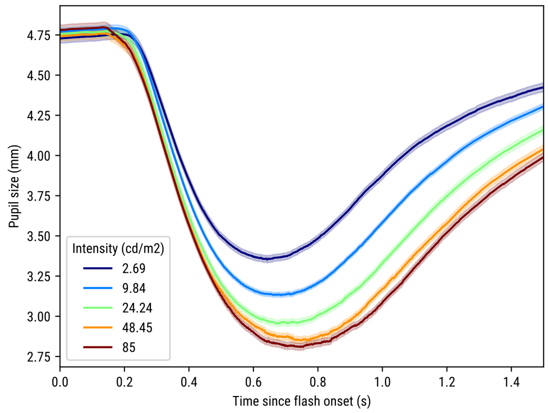


*Figure 1. Pupil size after flash onset as a function of stimulus intensity.*

### The first ERG component is not affected by blink artifacts

Participants frequently blinked. The first blink tended to occur around 200 ms after flash onset (Fig. 2). This was followed by a second and sustained period with blinks beginning around 500 ms after flash onset. Blink frequency varied considerably between participants with some participants blinking on only 5% of trials, while other participants blinked on more than 50% of trials (Fig. 2b).

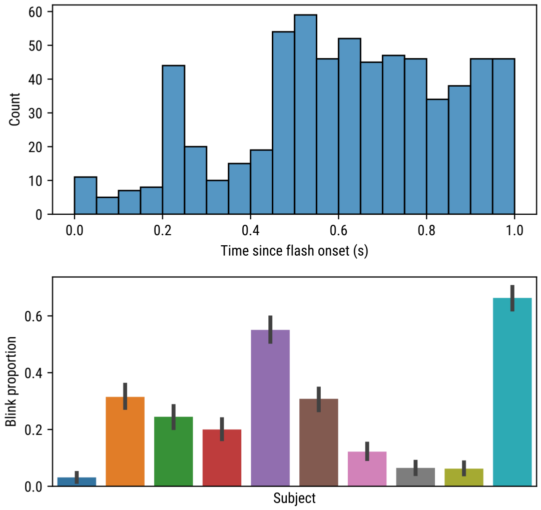


*Figure 2. a) Histogram of absolute blink frequency since flash onset for all participants and trials. b) Proportion of trials containing at least one blink for each participant.*

For the analyses presented in the main text, we excluded trials on which a blink occurred within 500 ms after flash onset. However, some blink artifacts likely remained, for example due to partial closures of the eyelid that were not detected as blinks by the eye tracker. During blinks, the eye lid increases its contact with the electrically positive cornea, allowing current to flow to the forehead (Matsuo et al., 1975; Picton et al., 2000). For this reason, blinks produce a positive potential at electrodes above the eyes but a negative potential at electrodes below the eyes. To investigate which parts of the ERG signal might be contaminated by blink artifacts, we therefore looked at upper- and lower-eyelid electrodes separately as a function of whether a blink was detected or not (before excluding any blinks; Fig. 3).

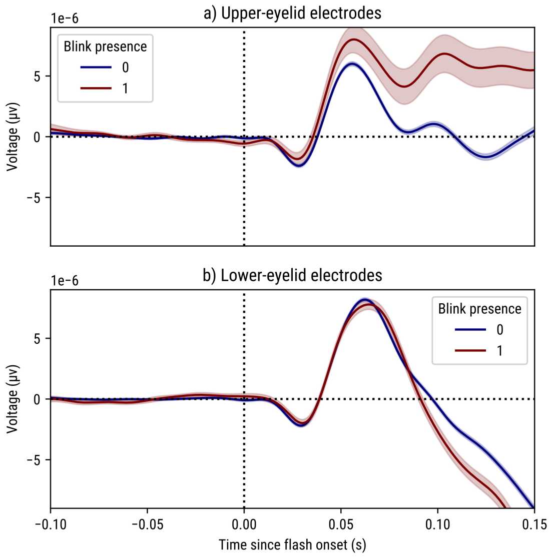


*Figure 3. a) Upper-eyelid electrodes as a function of blink presence since flash onset. b) Lower-eyelid electrodes as a function of blink presence since flash onset.*

Importantly, a visual inspection of the results reveals two key aspects of the data. First, the first component for the upper- and lower-eyelid electrodes is highly similar, indicating that it is not affected by blinks. However, after around 60 ms, the upper- and lower-eyelid electrodes start to diverge, and at the same time start to be modulated by the presence of blinks (as detected by the eye tracker); in other words, after around 60 ms, the ERG response is likely contaminated by blinks. For the upper eye-lid electrodes, the difference between blink and non-blink trials becomes apparent even earlier; visual inspection of individual trials (not shown) shows that this difference is driven by a handful of trials with very high voltages. (These trials were among the trials that were excluded for the main analyses.)

In addition, strong visual flashes can also induce a short-latency (e.g, 30-50 ms) visual startle blink reflex that is measurable even in the absence of actual eyelid movements as an electromyographic (EMG) signal from pairs of electrodes on the orbicularis oculi muscle (e.g., Bradley et al., 1990). Notably, however, such EMG activity was unlikely to contaminate our ERG measurements because the startle-EMG (1) should average out without prior rectification of the signals and (2) should be absent in our recording montage in which all eye electrodes are measured against a common reference (the mastoids) rather than against each other (as with a bipolar EMG reference).

Taken together, our analyses suggest that the early ERG component that is the main focus of our study is not affected by blink artifacts.

#### The ERG response is not affected by carry-over effects from the previous trial

The average interval between flashes was 3317 ms. This relatively short interstimulus interval makes it possible that the ERG response on the current trial is modulated by the intensity of the flash on the previous trial. This is an important consideration, because as reported in the main text, we found that ERG amplitude is modulated by whether the pupil is dilating or constricting at the moment that the stimulus is presented (see main text Fig. 2b). And this relationship could be driven by carry-over effects from previous trials; specifically, if the flash on the previous trial was of high intensity, this would cause a pronounced pupil constriction on the previous trial (see Fig. 1), and thus potentially a recovery in the form of pupil dilation on the current trial. In addition, if the flash on the previous trial was of high intensity, this could affect the ERG response on the current trial for example by altering the excitability of the retina. Via this route, carry-over effects could drive an association between pupil dilation and ERG amplitude.

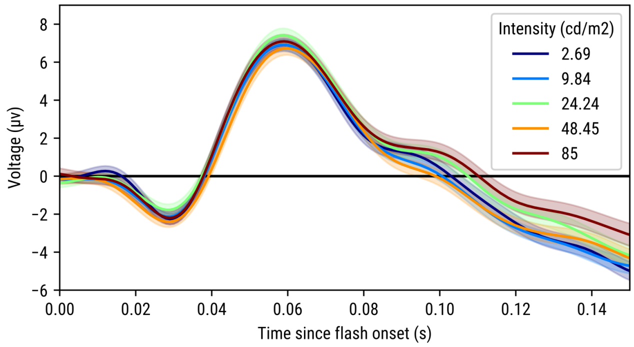


*Figure 4. ERG amplitude as a function of flash intensity on the previous trial.*

To test this, we selected trials on which a full-screen flash was presented and that were preceded by a (not previously excluded) trial on which a full-screen flash was also presented (on some trials, only the left, right, upper, or lower half of the display was used, see below); this left 2201 trials (47%) for analysis. Next, we analyzed whether the intensity of the previous trial affected the ERG response (Fig. 4), using the same model as discussed in the main text, except that the stimulus intensity of the current trial was replaced by that of the previous trial. This analysis revealed no significant effects of the intensity of the preceding flash on the ERG amplitude on the current trial.

Taken together, the ERG response is not notably affected by carry-over effects from the previous trial.

### Granger causality shows that ERG activity precedes and predicts occipital ERP activity

Retinal activity should precede and predict visual brain activity. To verify that this was indeed the case, we performed a Granger-causality analysis.

For each participant separately, we performed the following steps: 1) We determined the first derivative of the mean ERG and occipital ERP signal. Taking the derivative is a way to improve the stationarity of a signal (i.e. having a relatively constant mean and variance over time), which is an assumption of Granger-causality tests. 2) We calculated the F-value for the Granger-causality test (`statsmodels.tsa.stattools.grangercausalitytests`) from ERG to occipital ERP, and from occipital ERP to ERG; this was done for lags 10 to 30, where lag 10 means that the test considers all temporal lags from 0 ms to 10 ms in steps of 1 ms.

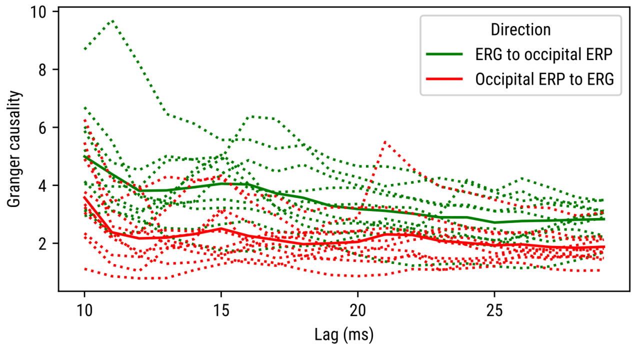


*Figure 6. Results of Granger causality tests. Dotted lines are individual subjects. Solid lines are mean values.*

Next, we performed a paired-samples t-test for the F-values from ERG to occipital ERP against the F-values from occipital ERP to ERG (Fig. 6). This was done separately for each lag. This revealed that there was more Granger causality from ERG to occipital ERP than the other way around for lags 10 to 20 (all $p < .05$).

Taken together, this indicates that ERG activity predicts and precedes occipital ERP activity.

#### Visual field manipulations affect ERG and ERP responses differently

While piloting the experiment, we were initially concerned that the signal picked up by the eye electrodes might originate from the brain (via volume conduction), rather than the retina. Based on the time course of the response and the topography of the early ERG response (see main text

Fig. 1c) we are now confident that this is not the case. However, this initial concern was the motivation for including trials in which a maximum intensity flash was presented only in the right, left, upper, or lower half of the display. The rationale behind this was that the effect of left-vs-right and upper-vs-lower visual field should be qualitatively different between ERG and occipital ERP responses. As shown below, this was indeed the case. We do not interpret these results beyond noting that, indeed, visual field manipulations have qualitatively different effects on ERG and occipital ERP responses.

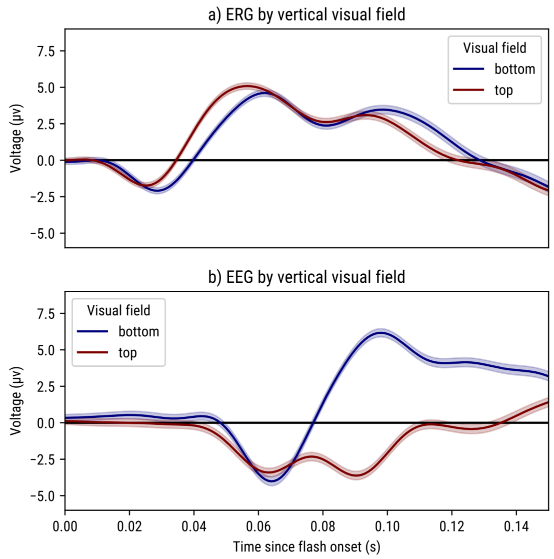


*Figure 7. The effect of vertical visual field presentation (bottom vs top) on a) ERG and b) occipital ERP amplitude (averaged across electrodes O1, Oz, O2).*

As shown in Fig. 7, the ERG response was slightly faster for upper as compared to lower visual field flashes but otherwise unaffected. Possibly, this reflects the monitor properties, such that the upper part of the display is refreshed ahead of the lower part. In contrast, and as expected, the occipital ERP response was strongly modulated by the stimulus position as of 70 ms after the flash, with flashes in the lower visual field eliciting stronger responses than flashes in the upper visual field (lower-hemifield advantage in visually evoked potentials, e.g. Capilla et al., 2016) (lower-hemifield advantage in visually-evoked potentials, e.g., Capilla et al., 2016).

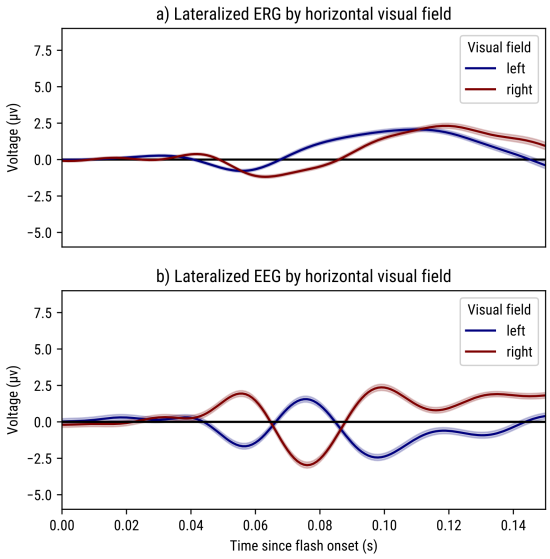


*Figure 8. The effect of horizontal visual field presentation (left vs right) on a) ERG and b) occipital ERP amplitude.*

To look at the effect of left- vs right-visual-field flashes, we analyzed the difference between left- and right-eye electrodes and left (O1)- and right (O2)-occipital electrodes. As shown in Fig. 8, both ERG and occipital ERP responses are modulated by visual field, but in a qualitatively different way.

### Adding interaction terms to the primary statistical model does not change the conclusions

To reduce model complexity, the statistical analysis in the main text is based on a model that includes stimulus intensity, pupil size, and pupil-size change as fixed effects, but does not include any interaction terms. A model with interaction terms, using ERG amplitude as dependent measure, yields very similar results. Notably, there were no clusters with interactions that spanned the earliest ERG component.

- Main effect of stimulus intensity: 12 - 36 ms: $p = .004$, 38 - 64 ms, $p = .001$, 94 -150 ms: $p < .001$
- Main effect of pupil size: 36 - 56 ms: $p = .034$, 64 - 150 ms, $p < .001$
- Main effect of pupil-size change: 16 - 150 ms, $p < .001$
- Stimulus intensity × pupil size: 44 - 70 ms, $p = .019$, 130 - 148 ms, $p = .04$
- Stimulus intensity × pupil-size change: no significant clusters
- Pupil size × pupil-size change: 70 - 106 ms, $p = .038$
- Stimulus intensity × pupil size × pupil-size change: no significant clusters

### Monitor properties

We used a photodiode to determine the point at which pixels reached 10% of maximum luminance after a display refresh, and used this to determine the locking time point (i.e. time point 0) for the ERG and EEG signals. As shown in Figure 8a, the locking time point coincided with the onset of the first pixels at the top left of the display. As shown in Figure 8b, the onset of pixels at the bottom right of the display occurred around 8 ms later.

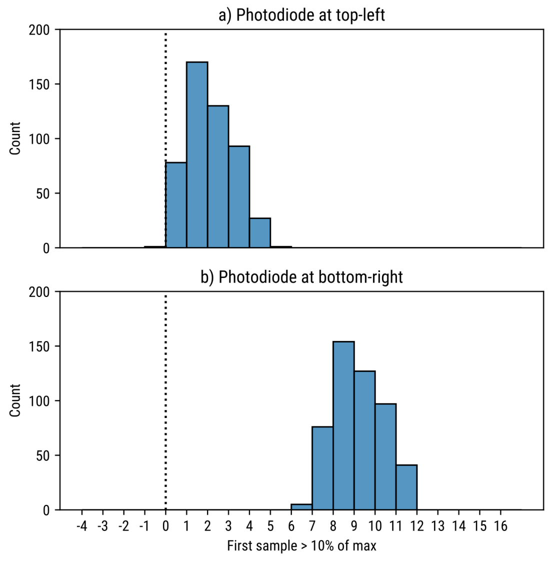


*Figure 9. a) Histogram of onsets (>10% of maximum luminance) of pixels on the top-left of the display relative to "time point 0", that is, the time point that we locked the ERG and EEG signals to. b) Histogram of onsets of pixels on the bottom-right of the display.*

The gradual filling of the display as well as the rise time of the pixels on the display (the time it takes for an individual pixel to reach maximum luminances; not shown) both affect the latencies of the ERG and EEG components reported in the main text.

### References


Capilla, A., Melcón, M., Kessel, D., Calderón, R., Pazo-Álvarez, P., & Carretié, L. (2016). Retinotopic mapping of visual event-related potentials. *Biological Psychology*, *118*, 114–125. https://doi.org/10.1016/j.biopsycho.2016.05.009

Matsuo, F., Peters, J. F., & Reilly, E. L. (1975). Electrical phenomena associated with movements of the eyelid. *Electroencephalography and Clinical Neurophysiology*, *38*(5), 507–511. https://doi.org/10.1016/0013-4694(75)90191-1

Picton, T. W., van Roon, P., Armilio, M. L., Berg, P., Ille, N., & Scherg, M. (2000). The correction of ocular artifacts: A topographic perspective. *Clinical Neurophysiology*, *111*(1), 53–65. https://doi.org/10.1016/S1388-2457(99)00227-8
